# Process-specific somatic mutation distributions vary with three-dimensional genome structure

**DOI:** 10.1101/426080

**Authors:** Kadir C. Akdemir, Victoria T. Le, Sarah Killcoyne, Devin A. King, Ya-Ping Li, Yanyan Tian, Akira Inoue, Samir Amin, Frederick S. Robinson, Rafael E. Herrera, Erica J. Lynn, Kin Chan, Sahil Seth, Leszek J. Klimczak, Moritz Gerstung, Dmitry A. Gordenin, John O’Brien, Lei Li, Roel G. Verhaak, Peter Campbell, Rebecca Fitzgerald, Ashby J. Morrison, Jesse R. Dixon, P. Andrew Futreal

## Abstract

Somatic mutations arise during the life history of a cell. Mutations occurring in cancer driver genes may ultimately lead to the development of clinically detectable disease. Nascent cancer lineages continue to acquire somatic mutations throughout the neoplastic process and during cancer evolution (Martincorena and Campbell, 2015). Extrinsic and endogenous mutagenic factors contribute to the accumulation of these somatic mutations (Zhang and Pellman, 2015). Understanding the underlying factors generating somatic mutations is crucial for developing potential preventive, therapeutic and clinical decisions. Earlier studies have revealed that DNA replication timing (Stamatoyannopoulos et al., 2009) and chromatin modifications (Schuster-Böckler and Lehner, 2012) are associated with variations in mutational density. What is unclear from these early studies, however, is whether all extrinsic and exogenous factors that drive somatic mutational processes share a similar relationship with chromatin state and structure. In order to understand the interplay between spatial genome organization and specific individual mutational processes, we report here a study of 3000 tumor-normal pair whole genome datasets from more than 40 different human cancer types. Our analyses revealed that different mutational processes lead to distinct somatic mutation distributions between chromatin folding domains. APOBEC- or MSI-related mutations are enriched in transcriptionally-active domains while mutations occurring due to tobacco-smoke, ultraviolet (UV) light exposure or a signature of unknown aetiology (signature 17) enrich predominantly in transcriptionally-inactive domains. Active mutational processes dictate the mutation distributions in cancer genomes, and we show that mutational distributions shift during cancer evolution upon mutational processes switch. Moreover, a dramatic instance of extreme chromatin structure in humans, that of the unique folding pattern of the inactive X-chromosome leads to distinct somatic mutation distribution on X chromosome in females compared to males in various cancer types. Overall, the interplay between three-dimensional genome organization and active mutational processes has a substantial influence on the large-scale mutation rate variations observed in human cancer.

---

The distribution of somatic mutations shows significant variation among cancer genomes, which is associated with the transcription binding sites (Perera et al., 2016; Sabarinathan et al., 2016), chromatin modifications (Makova and Hardison, 2015) and proximity to nuclear periphery (Smith et al., 2017). Three-dimensional genome structure is closely related to DNA replication and transcription (Pope et al., 2014). Chromatin conformation (Hi-C) studies have revealed cell-type invariant topologically associating domains (TADs) as an important component of genome-folding architecture (Dekker and Heard, 2015; Dixon et al., 2016). Contact frequencies of the regions within a TAD are higher compared to contact frequencies between neighboring TADs, and genes within the same TAD can exhibit coordinated expression patterns (Dixon et al., 2012; Fudenberg et al., 2016; Nora et al., 2012; Rao et al., 2014). However, whether there is a connection between this chromatin folding organization and somatic mutation rates in cancer genomes remains unclear. Diverse mutational processes lead to somatic mutations in human cancer (Martincorena and Campbell, 2015) and mutational signatures are used to delineate the imprints of underlying processes on cancer genomes (Alexandrov et al., 2013; Nik-Zainal et al., 2012). Understanding the interplay between chromatin folding and mutational signatures is important to elucidate the mechanisms behind different DNA repair and damage processes. Here, we sought to understand the relationship between genome organization and mutational processes observed from more than 60 million somatic mutations identified in whole genome sequencing datasets of 42 different histology subtypes (Supplementary Table 1).

## The distribution of somatic mutations and DNA repair (NER) activity is correlated with the three-dimensional genome organization

In order to evaluate the rate of somatic mutation accumulation in TADs, we annotated TADs in to five distinct groups, in which each domain was classified based on the most common chromatin state (histone-tail modifications) in the Roadmap Epigenome project cell types (methods). Each annotation group is enriched in different chromatin states (Supplementary Fig. 1a). We specifically focused on two domains types, namely transcriptionally-inactive (Low) and transcriptionally-active (Low-Active) because these domains cover 71% of the human genome (Supplementary Fig 1b). In agreement with previous studies (Lawrence et al., 2013; Stamatoyannopoulos et al., 2009), the mutation load of a given domain is negatively correlated with its average replication timing (r^2^ between −0.48 to −.68; Supplementary Fig. 1c). Notably, mutation load and chromatin state differed drastically across the TAD boundaries if neighboring TADs are epigenetically distinct (Fig. 1a, Supplementary Fig. 1d), which implies that the distribution of somatic mutations in cancer genomes corresponds to spatial chromatin organization. This observation prompted us to investigate the average distribution of mutations on either side of TAD boundaries between epigenetically distinct and similar TADs. We found that average mutation load is drastically different around the boundaries between epigenetically distinct TADs compared to the boundaries separating transcriptionally similar TADs (Fig. 1b, Supplementary Fig. 1e). In addition, we observed a depletion of somatic mutations at TAD boundaries compared to flanking regions which can be attributed to higher transcriptionally-activity at TAD boundaries (Fig. 1c).

**Figure 1.**
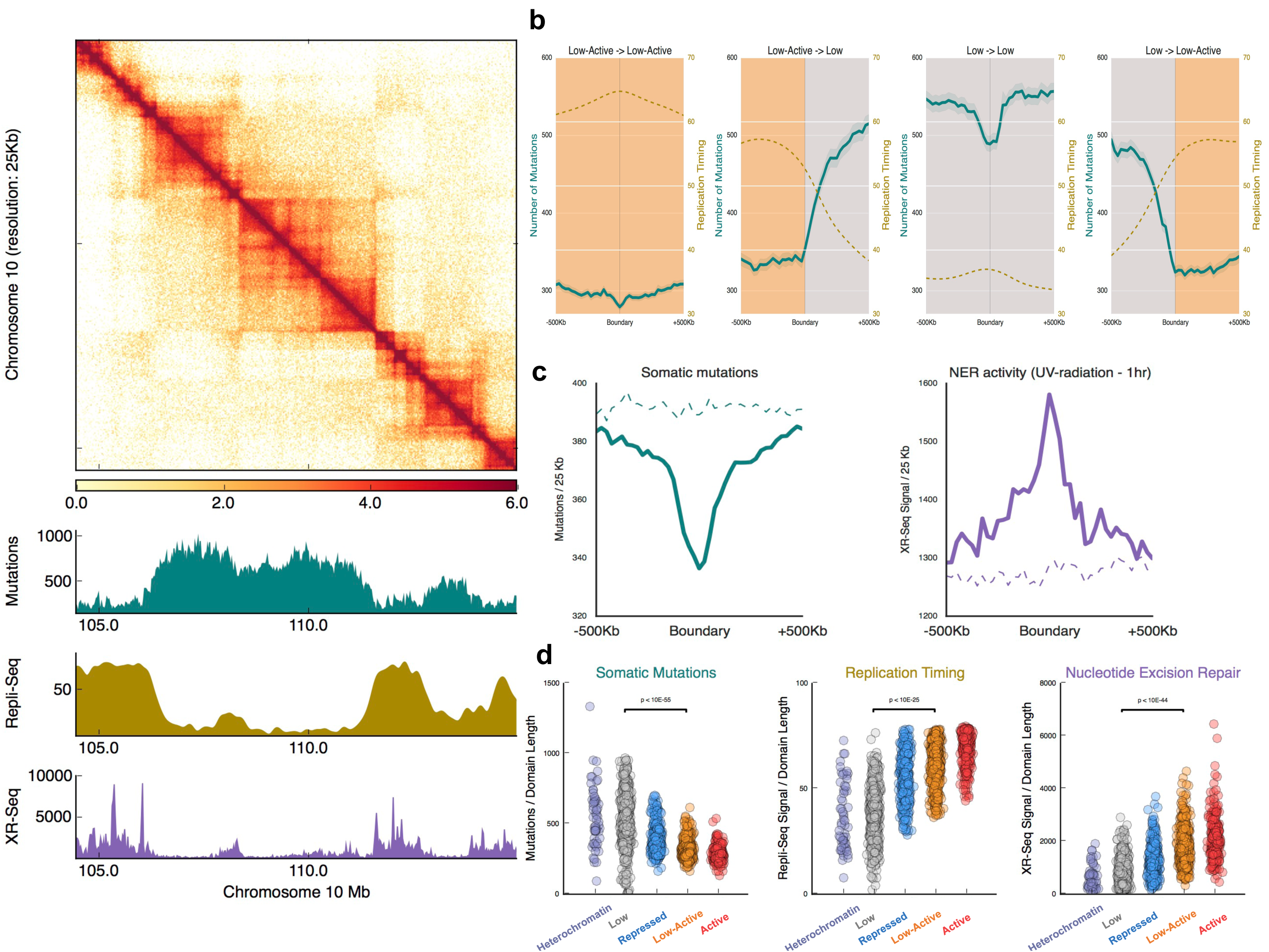
The distribution of somatic mutations in cancer genomes and NER activity is correlated with the three-dimensional genome organization. **a)** Contact matrix shows the Hi-C chromatin contact map from keratinocyte cells (NHEK). Below, histograms show the distributions of somatic mutation accumulation in 3000 cancer samples (green), average replication timing (gold) and chromatin binding position of nucleotide-excision repair upon UV damage in IMR90 fibroblast cells (purple) for the corresponding genomic region. **b)** Mutational load is correlated with spatial chromatin organization. Average profiles of somatic mutation accumulation in 3000 cancer samples (green), average replication timing (gold) across 500Kb of boundaries delineating: low-active (orange) to low-active (orange); low-active (orange) to low (gray); low (gray) to low (gray); and low (gray) to low-active (orange) TADs. **c)** Profiles of somatic mutations and NER enrichments around TAD boundaries. Dashed lines represent the enrichment profiles at shuffled boundaries. **d)** Dot plots show the significant difference in accumulation of somatic mutations, average replication timing and NER binding between transcriptionally-inactive TADs (Low) and transcriptionally-active TADs (Low-active). Enrichment of somatic mutations, replication timing and NER binding are normalized with the domain length.

These results highlight an implicit link between spatial chromatin organization and regional mutation rates in cancer cells. What is unclear, however, is whether this reflects an increased mutational burden in inactive regions of the genome, or whether the differences are the result of differential ability to repair somatic mutations depending on chromatin state. To assess the potential interplay of chromatin folding, DNA repair activity and mutational distribution, we used nucleotide-excision repair (NER) DNA binding profiles (XR-Seq) in UV-exposed human skin fibroblast cells (Hu et al., 2015). NER complexes demonstrate a distribution pattern that was correlated with TAD architecture (Fig 1a). However, in contrast to somatic mutation rates, transcriptionally-active TADs (low-active) are significantly enriched in NER activity compared to transcriptionally-inactive TADs (low) (Fig. 1d). We also utilized UV-mediated DNA-lesion (cyclobutane pyrimidine dimers) locations in human fibroblast cells (García-Nieto et al., 2017) and observed that inactive TADs acquire significantly higher damage compared to active TADs (Supplementary Fig. 1f). Therefore, the combination of higher localization of NER in transcriptionally-active regions (including the boundaries) and higher rates of DNA damage lesions in transcriptionally-inactive domains partially explains the lower mutational load in active regions compared to inactive regions, at least for these specific classes of mutagens. Given that the NER complex is coupled with transcriptional apparatus (TC-NER), the higher enrichment of NER activity in transcriptionally-active domains overlaps with earlier findings (Kamarthapu and Nudler, 2015). However, at later time points of the UV damage, NER enrichment becomes higher in inactive TADs compared to active TADs (Supplementary Fig. 1g), suggesting a shift in DNA repair activity at different time points upon DNA damage. Our results imply that the distribution of somatic mutations and DNA repair activity is closely related with the three-dimensional genome organization.

## Hypermutated tumors exhibit variation in regional mutation rate between transcriptionally–active and –inactive TADs

Distinct mutational processes are active and contribute to overall somatic mutation distribution in a given tumor. Therefore, we investigated how individual processes generate mutational distributions. First, we focused on hyper-mutated cancers that can be associated with different underlying mutational processes. We classified samples as hypermutated if they harbor more than 66 somatic mutations per megabase, compared to the average of ~6.6 mutations per megabase in all samples (Supplementary Fig. 2a-b, Fig. 2a). High levels of somatic mutations could be due to DNA-repair deficiencies and/or exposure to potent mutagens. To evaluate these different processes, we initially focused on specific samples driven by diverse mutational processes. Specifically, we analyzed three microsatellite-instability (MSI) cases, where DNA mismatch repair (MMR) activity is deficient due to a) inherited biallelic mutations (PMS2 c.1866G>A/c.1840A>T) in a case of childhood glioma, b) MLH1 silenced by DNA methylation in a case of uterine cancer, c) repair deficient adult glioma, induced by a DNA-alkylating agent (temozolomide) (Kim et al., 2015) (MSH6 c.97G>A/c.289G>A/c.1058G>A). In addition, a colon cancer sample where mutations in proofreading polymerase (PoLε c.3675A>G) resulted in high numbers of mutations was examined. Furthermore, we examined cases where exogenous genotoxin exposure can lead to hypermutation such as UV-exposed melanoma samples (Supplementary Fig. 2a) and a case of urinary tract urothelial carcinoma (UC) where a potent mutagen, aristolohic acid (AA), causes hypermutations. UV-mediated skin cancer, PoLε-deficient colon cancer and AA-induced urinary tract UC exhibit 2.6-, 1.75-, 1.5-fold higher mutation rate in inactive domains compared to the active domains, respectively. On the other hand, all of the MSI samples, despite having different mutational signature, have a similar mutation burden between transcriptionally–active and –inactive TADs (flatter mutation distribution, Fig. 2b-d). Both inherited (PoLε c.2237G>T/c.564G>T/c.461A>G) and sporadic MSI sample (PoLε c.465C>T) also harbor somatically-acquired Polε-deficiency but unlike the sample with only PoLε-deficiency (PoLε c.3675A>G), these samples demonstrate a flatter mutation distribution. We extended our analysis to all samples and noticed that, in case of concomitant MSI and Polε-deficiencies, flatter mutational distribution is observed around chromatin folding domains (Supplementary Fig. 2c). This observation is in agreement with an earlier study describing a decreased mutation load variation in MSI and Polε-deficient tumor genomes (Supek and Lehner, 2015). NER is the main DNA repair pathway to repair AA-induced A>T mutations (signature 22) and UV-mediated predominantly C>T mutations (signature 7). However, we observed a difference between the distributions of mutations across TADs for these two mutagens, where the relative rate of mutations in active domains is higher for AA-induced mutations compared to UV-related mutations (Fig. 2b). This observation implies a potential mechanistic and potency difference between these mutagens, and studying the distribution of mutations may reflect the nature of differences in DNA repair upon the exposure to distinct mutagens.

**Figure 2.**
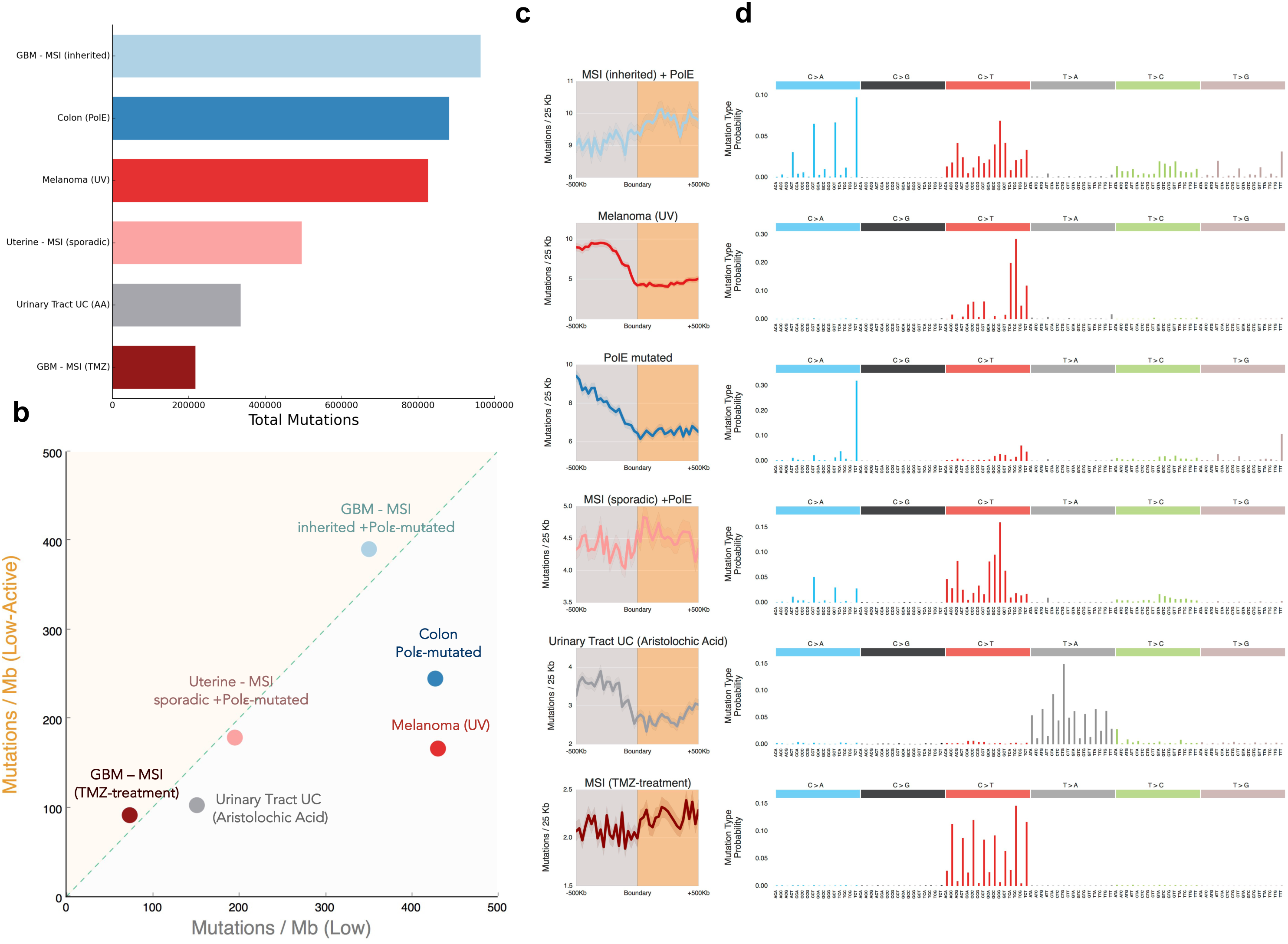
Hypermutated tumors exhibit variation in regional mutation rate between transcriptionally –active and –inactive TADs. **a)** Bar plot shows the number of somatic mutations per hypermutated sample. **b)** Number of mutations per megabase in transcriptionally–active (Low-Active) TADs versus transcriptionally-inactive (Low) TADs. **c)** Aggregate plot shows the somatic mutations around TAD boundaries delineating transcriptionally-inactive (Low) TADs from transcriptionally-active (Low-active) TADs. **d)** Mutational signature for each sample. The probability bars for the six types of substitutions are displayed in different colors.

## Higher order chromatin organization in colorectal cancer cells

We next explored if the flatter mutation distribution in hypermutated MSI cases studied initially could be observed in MSI samples of various cancer types. To this end, we extended our analysis to all samples identified as MSI (methods), and indeed, the flatter mutation distribution pattern is apparent in MSI samples of different cancer types (Supplementary Fig 3a-b). For example, MSI samples identified in colon (8 samples) or gastric (10 samples) adenocarcinoma cohorts exhibit the flatter mutation distributions around TAD boundaries where a distinct difference is observed in DNA mismatch repair proficient (MSS) samples (Fig. 3a). This flatter distribution is not dependent on the mutational load as we observed a low mutation burden colon MSI sample with a similar flatter mutation distribution (Supplementary Fig. 3c). One possibility is that the difference in mutational distribution is to due altered 3D genome structure in MSI samples. In order to understand whether this flatter mutational distribution is a result of higher-order chromatin reorganization in MSI cells, we generated high-resolution Hi-C datasets on two DNA-repair proficient (SW480, CaCo2) and two DNA mismatch-repair deficient (LoVo, DLD1) colon cancer cell lines. LoVo cell line has MMR-deficiency (MSH3: c.2384T>A), whereas DLD1 cell line contains both DNA Polymerase proofreading-(PolD1 c.29G>T/c.2065C>T/c.2237G>T) and MMR-(MSH3: c.1463T>C; MSH6: c.1406A>T) deficiencies. DNA-repair status of these cell lines can be also observed from their mutational signature profiles (Fig. 3b). As a reference for non-malignant colon cells, we reprocessed Hi-C data from a healthy colon tissue sample (GTEx Consortium, 2013). We first focused on chromatin interactions around a boundary between an active and an inactive TAD where a stark difference on mutation load was observed in MSS and MSI samples. Interestingly, chromatin folding patterns are similar around this boundary in MSS and MSI cancer cell lines despite the flatter mutational distribution in MSI samples (Fig. 3b). We next profiled chromatin interactions across all TAD boundaries and noticed that TAD boundary strengths of MSI cancer cells are comparable to that of healthy colon and MSS cancer cells (Fig. 3c-d). Having ruled out that MSI in and of itself was a primary effector of chromatin folding, we sought to further elucidate the underpinnings of flatter distribution in MSI samples, given that the overall chromatin architecture remains unaffected. Short tandem repeats (STRs) are most prone to acquire new mutations in case of DNA mismatch-repair deficiency (Kunkel and Erie, 2015). We profiled the distribution of STRs in the human genome (Gymrek et al., 2017) and noticed that active TADs harbor significantly higher repeat content (Fig. 3e, Supplementary Fig. 3d). Repeat region mutations in MSI tumors are significantly enriched in active domains (Fig. 3e), whereas non-repeat site mutations do not exhibit any enrichment in active domains (Supplementary Fig. 3e). On the contrary, MSS tumor mutations are enriched at inactive domains for both repeat and non-repeat sites (Fig. 3e, Supplementary Fig. 3e). Therefore, flatter mutation load in MSI samples can be attributed to an interplay between the uneven distribution of repeat contends and differential repair activity across TADs.

**Figure 3.**
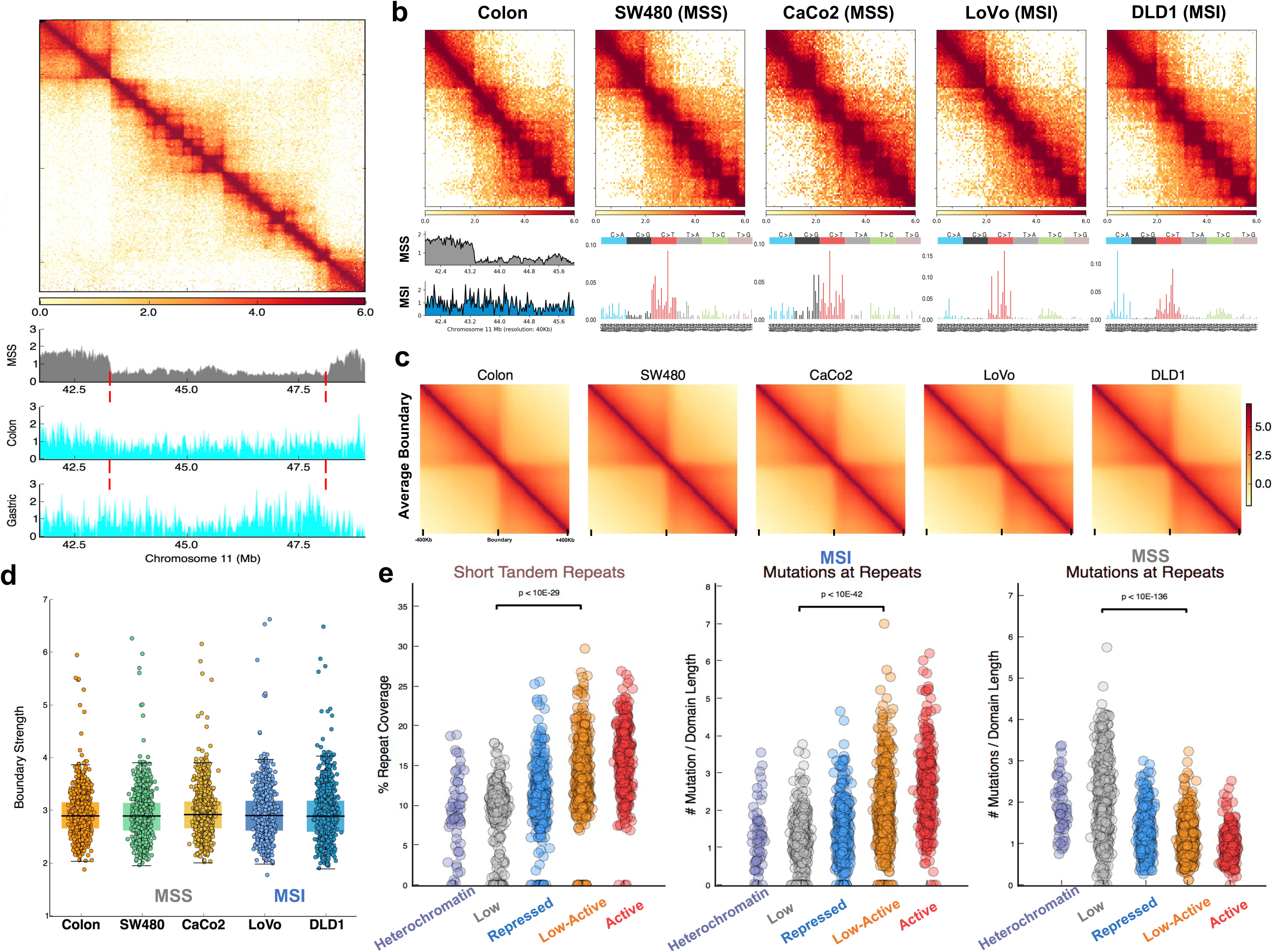
Higher order chromatin organization in DNA mismatch repair deficient and proficient cancer cells. **a)** An example region represents the mutation distribution differences between MSS (gray) and MSI (aqua) cancer samples with respect to chromatin folding. Histograms represent normalized mutation counts for MSS and MSI samples. Normalization is performed by dividing the sum of mutations in a cohort to hundred thousand. Contact matrix shows the Hi-C chromatin contact map from keratinocyte cells (NHEK). Red dashed-bars represent boundaries between transcriptionally inactive and active TADs. **b)** Chromatin interaction maps of healthy colon tissue and colon cancer cell lines: DNA-repair proficient SW480 and CaCo2, MSI: LoVo, MSI and Polymerase deficiency: DLD1 around the boundary between inactive and active domains represented in figure 3a Bottom panel shows mutational signature for cancer cell line and distribution of somatic mutations in MSS and MSI colon cancers (left). **c)** Average contact enrichment and d) average TAD boundary strength profiles across all TAD boundaries in healthy colon tissue and colon cancer cell line Hi-C data. Scored used described in Supplementary Methods. **e)** Distribution of short tandem repeats in different domain types (left). Mutations at STR sites are significantly enriched at active domains in MSI tumors (middle), whereas repeat region mutations are enriched at inactive domains in MSS tumors (right).

## Mutation distribution depends on the active mutational processes

Multiple mutational processes could be active in a tumor such as PoLε-deficiency in the context of MMR deficient tumors (Fig. 2b). We further sought to identify cases where different mutational processes are operative within an MSI sample that might change the flatter somatic mutations. Our analysis demonstrated that signature 17, most commonly observed in gastric and esophageal adenocarcinoma, exhibits a high mutation preference for inactive domains (Supplementary Table 2). A recent study reported the occurrence of MMR deficiency, albeit in low frequency, in esophageal adenocarcinoma (Secrier et al., 2016). Therefore, we examined the mutation distributions in three MSI esophageal adenocarcinoma samples (labeled as 5935, H03, E03) with co-occurrence of signature 6 (MSI-related) and signature 17. Signature 6 enrichment (indicative of MMR deficiency) is at similar levels (5935: 0.37, H03: 0.30, E03: 0.35; these ratios represent weights of signatures calculated by deconstructSigs (Rosenthal et al., 2016) algorithm) in all three samples. Distribution of somatic mutations with respect to TADs revealed that the MSI esophageal sample with only signature 6 enrichment exhibits flat mutation rate as expected in MSI cases (5935), whereas MSS esophageal samples with only signature 17 enrichment (gray dots) exhibit a high mutation burden in inactive domains. An MSS sample (A920) is shown in figure 4a, indicated by a black dot as an example. However, MSI samples with signature 17 enrichments (H03: 0.48 and E02: 0.14) deviate from the flatter distribution and show a slightly higher mutational load in inactive TADs (Fig. 4a). The degree of change in mutation distributions around TADs is correlated with the contribution of signature 17 (Supplementary Fig. 4a). Signature 6 compromises mutations mainly with C>T transition whereas signature 17 mutations are predominantly T>G mutations (Supplementary Fig. 4a-b). Indeed, the active domain mutations in MSI samples are positively correlated with the higher ratio of C>T mutations (r^2^: 0.96) compared to negative correlation observed in the T>G mutations (r^2^: −0.99) (Supplementary Fig. 4c-d).

**Figure 4.**
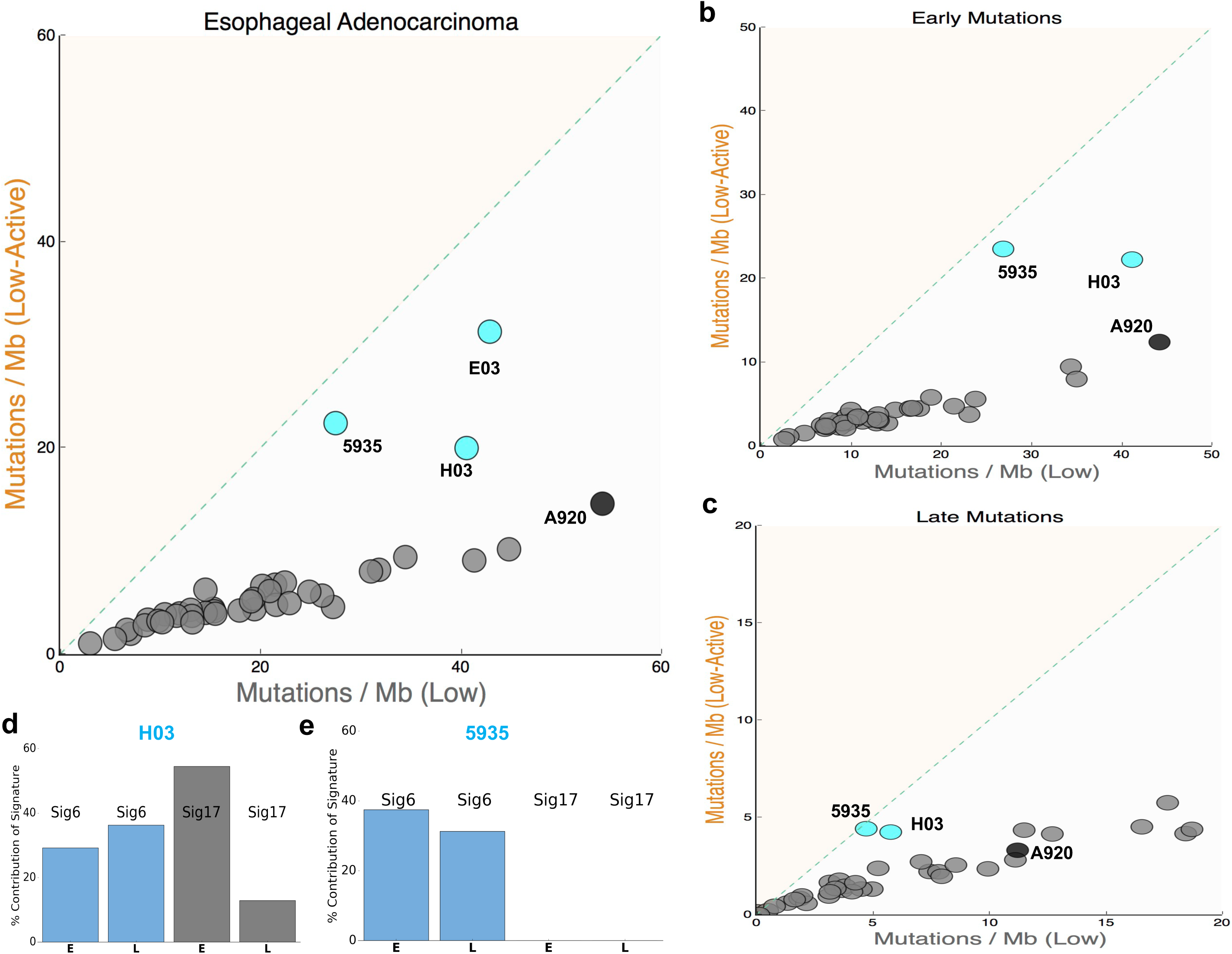
Mutation distribution depends on the active mutational processes within a tumor and subjects to changes during tumor evolution. **a)** Number of mutations per megabase in transcriptionally–active TADs (Low-active) versus transcriptionally-inactive TADs (Low). Esophageal adenocarcinoma samples with MMR-deficiency (aqua dots) show various mutation distributions based on signature 17 enrichment levels. Gray dots represent signature 17 enriched esophageal adenocarcinoma samples. **b-c)** Number of mutations per megabase in transcriptionally–active TADs (Low-active) versus transcriptionally-inactive TADs (Low) for **b)** early and **c)** late mutations. Esophageal adenocarcinoma samples with MMR-deficiency denoted with aqua dots, gray dots represent signature 17 enriched esophageal adenocarcinoma samples. **d-e)** Bar plots shows the percent contribution (calculated by deconstructSigs algorithm) of signature 6 (aqua) and signature 17 (gray) in early (E) and late (L) mutation burden **d)** H03 and **e)** 5935 samples.

Mutational processes might be active at different stages of tumor evolution. Therefore, we sought to characterize the relative timing of signature 6 and signature 17 activities during tumor evolution. Through interrogation of allelic fraction of each mutation (methods), we were able to attribute the timing of mutations as early or late for two MSI samples (5935 and H03) and rest of the MSS esophageal adenocarcinoma samples (Supplementary Fig. 4e). We noticed that signature 6 and 17 could be detected at the early stage mutations in H03 sample, thus the distribution of mutations deviate from MSI-like flatter profile (Fig. 4b). However, in late mutations of H03 sample, signature 6 level remains similar, while signature 17 enrichment is lower (Fig. 4c-e), which can be observed in the ratio of T>G mutations (Supplementary Fig. 4f-g). This results in the distribution of mutations being different between early and late mutations (early 1.84-fold, late 1.06-fold higher mutations in inactive domains compared to active domains) in H03 where late mutations exhibit a flatter distribution (Fig. 4b-c). On the other hand, 5935 and A920 signatures remain similar for early and late mutations (Fig. 4e, Supplementary Fig. 4f-g), with the distribution of mutations remaining unchanged for these samples (Fig. 4b-c). Overall, when multiple mutational processes are active in a tumor sample, timing of distinct processes affect the global distribution of somatic mutations along the genome.

## APOBEC-related mutations are enriched in transcriptionally-active TADs

To identify additional mutational processes causing the differential distributions across the chromatin folding domains, we calculated the slope of mutations in samples as the ratio of mutation rates in transcriptionally-active and –inactive TADs (methods). We noted that samples with high signatures 2 and 13 enrichments exhibit a positive correlation (r^2^: 0.40) with the slope of mutations suggesting a higher preference in active domains compared to inactive domains (Fig. 5a and Supplementary Fig. 5a). These signatures are attributed to the aberrant activity of APOBEC family deaminases. We further assessed the relationship between the slope of mutations and fold enrichment of stringent APOBEC-related mutation motif (tCa>tTa or tCa>tGa; henceforth referred as tCa) over random mutagenesis expectation. We observed that samples with higher APOBEC-mutagenesis fold enrichment exhibit a higher correlation with the slope of mutations (high-enrichment r^2^: 0.58; Not-APOBEC r^2^: 0), consistent with earlier studies about APOBEC mutagenesis enrichment in early replicating regions (Kazanov et al., 2015). APOBEC family enzymes have been implicated in the genesis of localized hypermutations (kataegis loci), based on their biochemical specificity to single-strand (ss) DNA, specifically in the context of tCw>tGw or tCw>tTw (W= A or T, mutated base capitalized) mutations (Fig. 5c). We therefore profiled the distributions of the kataegis-like loci (C- or G-coordinated mutation clusters) with respect to three-dimensional genome architecture. Kataegis-like loci significantly overlap with TAD boundaries and TADs with higher transcriptional activity (p-values < 10^-5^). However, these loci are significantly depleted at transcriptionally-inactive TADs (low TADs; p-value < 10^-5^) (Fig. 5d, Supplementary Fig. 5b). Previous studies reported that clustered APOBEC-mutagenesis occur in ssDNA during replication or in association with repair of double-strand breaks (Chan and Gordenin, 2015). Our observation suggests that chromatin folding features could affect the distribution of persistent ssDNA prone to hypermutation.

**Figure 5.**
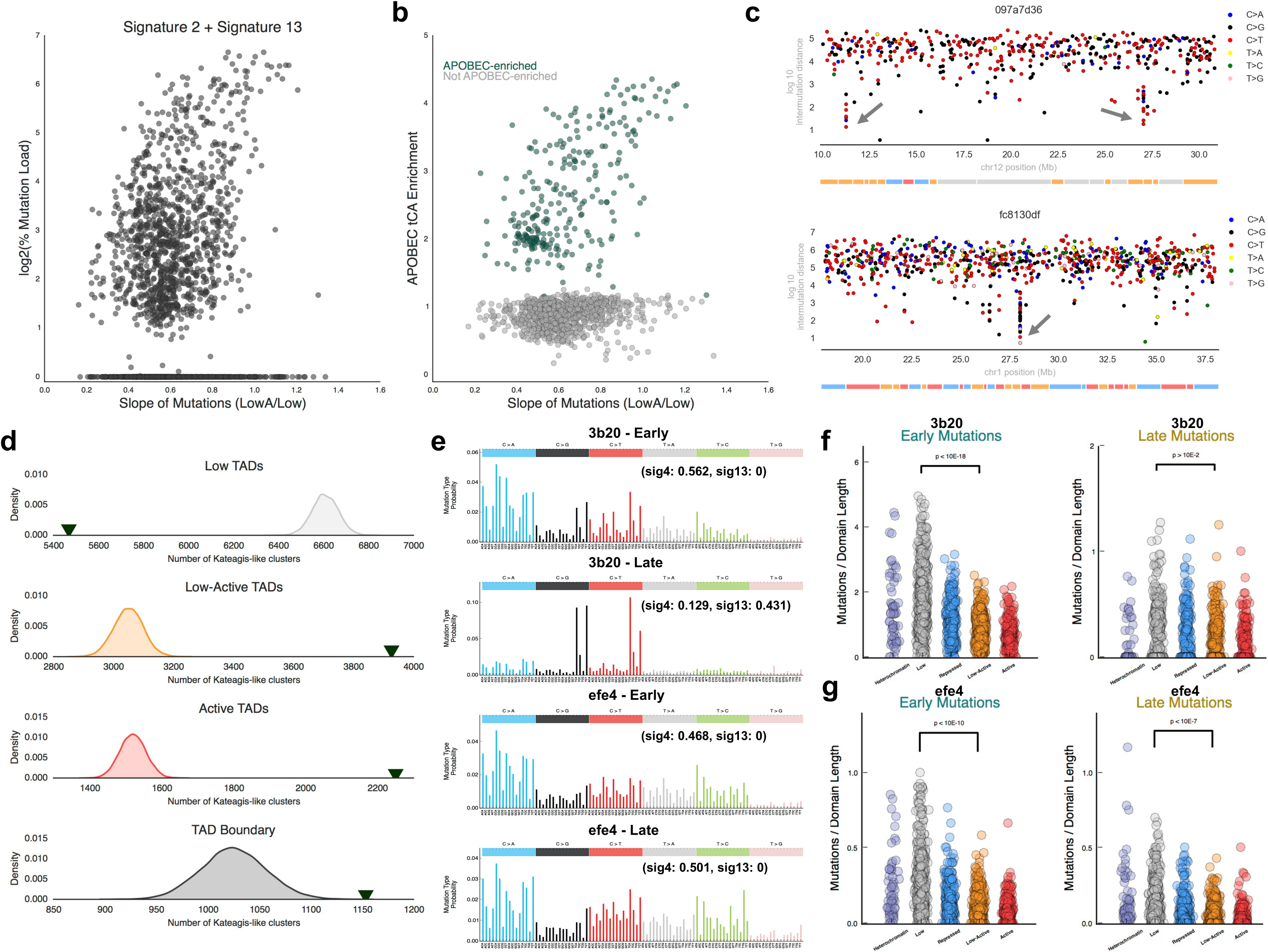
APOBEC-related mutations are enriched in transcriptionally-active TADs. **a)** Association between APOBEC-related mutation signatures (sum of signatures 2 and 13) enrichment and slope of mutations (LowA: transcriptionally-active TADs versus Low: transcriptionally-inactive TADs). **b)** The distribution of APOBEC-signature enrichment subtypes (dark-green: A3A-like, gray: Not APOBEC-enriched) versus slope of mutations (LowA: transcriptionally-active TADs versus Low: transcriptionally-inactive TADs). **c)** APOBEC-related clustered mutations (kataegis) favor transcriptionally active TADs. Kataegis-like clusters were identified as described in Chan et al. Example kataegis loci is shown with log10 intermutation distance on y-axis in two different breast cancer samples. Each dot represents a mutation; colors indicate different base substitution. Rectangular boxes below depict individual TADs within the given genomic locus (gray and blue colors for transcriptionally-inactive TADs; orange and red colors show transcriptionally-active TADs). **d)** Dark green arrows represent overlap of kataegis loci with different TAD-types and – boundaries. Curves indicate the distribution of the expected overlap based on randomized data. Mutational signature for early and late mutations of lung cancer samples (3d20: top; efe4: bottom). Late mutations change to APOBEC signature in 3d20, whereas remains as tobacco signature in efe4 sample. The probability bars for the six types of substitutions are displayed in different colors. Contributions of each signature to observed mutation load are denoted on upper left corner. **f-g)** Dot plots show the difference in accumulation of early and late somatic mutation in sample **f)** 3d20 accumulations and **g)** similar profiles in sample efe4 between transcriptionally-inactive TADs (gray) and transcriptionally-active TADs (orange). Enrichment of somatic mutations are normalized with the domain length.

Tobacco smoke has been postulated to indirectly activate APOBEC-related mutagenesis in certain cancer types (Alexandrov et al., 2016). We focused on the timing of mutations in lung cancers where tobacco smoke carcinogens are the main etiologic agent of the disease. Mutational signature 4, which is strongly correlated with smoking (Alexandrov et al., 2016), and has the strongest enrichment in the lung cancer samples. We noticed that in certain tumor samples the early mutations are enriched in signature 4 but late stage mutations reflect APOBEC signatures indicating a shift of active mutational processes during somatic evolution of cancers (Fig. 5e). For example, in sample 3d20, early mutations are significantly enriched in inactive domains (p-value < 10^-17^, Fig. 5f) but late mutations, exhibiting APOBEC signature, are distributed more evenly between domains (p-value > 10^-2^, Fig. 5f). On the other hand, in sample efe4, signature 4 is persistent throughout the tumor development (Fig. 5e). Consequently, both early (p-value < 10^-10^) and late (p-value < 10^-7^) mutations are significantly enriched in inactive domains (Fig. 5g). Overall, the samples with early smoking signature and late APOBEC signature during tumor progression exhibit significant (p-value: 0.009) change in mutation distributions (Supplementary Fig. 5c). In contrast, samples with constant tobacco signature enrichment during early and late stages of the tumor development do not show any significant change (Supplementary Fig. 5c). These results highlight that mutational distributions in cancer genomes can shift during tumor evolution based on the active mutational processes. In addition to tobacco smoking activation, APOBEC enzymes can be activated upon virus infection as this family of enzymes constitutes parts of the innate immune response against viruses (Chan and Gordenin, 2015), Hence, we profiled mutational distributions in virus related cancer types. Human papillomavirus (HPV)-positive head and neck squamous cell (HNSC) carcinoma samples exhibit higher APOBEC enzyme expression and have a significantly higher tendency to accumulate mutations in active TADs compared to HPV-negative HNSC samples (Supplementary Fig. 5d). However, we did not notice any enrichment for APOBEC mutation signatures in Hepatitis B virus (HBV) and Hepatitis C virus (HCV) associated liver cancers or Epstein-barr virus (EBV) associated gastric cancers (Supplementary Fig. 5f). Therefore, APOBEC-mediated mutational load distribution between TADs is specific to certain virus types and is observed more frequently in HPV-infected HNSC cancers.

Taken together, active mutational processes dictate the mutation distributions across the genome in distinct stages of cancer evolution and bulk tumor sequencing reflects the composite genome distribution patterns from various mutational processes over time.

## Patterns of different mutational signatures

In addition to APOBEC- and MSI-associated signatures (Supplementary Fig. 5a and 6a), we identified several additional mutational signatures exhibiting distinct accumulation patterns along chromatin folding domains (Supplementary Fig. 6). For example, mutational signature (signature 3) associated with the failure of DNA double-strand break-repair by homologous recombination, exhibits a higher preference toward active domains (Supplementary Fig. 6a). Two signatures associated with base excision repair deficiency (signature 30 and signature 36) also exhibit a positive correlation with the active domains (Supplementary Fig. 6a). Overall, we noticed that the known exogenous damage agents such as UV-light (signature 7), tobacco-smoke (signature 4) or acid reflux-related (signature 17) generates more mutations in inactive domains, whereas endogenous factors such as APOBEC, MMR- or base excision repair-deficiency mutations are enriched in active domains (Supplementary Fig. 6a-b). Increased mutation load in active domains in DNA repair deficiency cases is overlapping with the findings of an earlier study which reported that NER deficient melanoma samples have a higher mutation burden in open chromatin regions compared to NER proficient melanomas (Polak et al., 2014). Polymerase deficiencies are an exception to this dichotomy. Although endogenous factors; both PoLε (signature 10) and PoLh (signature 9) deficiencies generate mutations in inactive domains (Supplementary Fig. 6c-d). In addition, a positive correlation between mutational signature enrichment and active domain preferences are observed for mutational signatures of unknown aetiology, such as signature 16 and 28 (Supplementary Fig. 6e-f). Interestingly, signature 28 co-occurs with Polε-deficiency in uterus and colorectal adenocarcinoma samples and signature 28 mutations are predominantly T>G, similar to signature 17 mutations, but unlike signature 17 distribution, signature 28 mutations generally occur in the active domains (Supplementary Fig. 6f-g). This observation indicates that different mutational signatures with overlapping sequence context could generate distinct mutational distribution. Similarly, signatures 12 and 16 are mostly observed in liver cancers with a strong transcriptional strand bias (Haradhvala et al., 2016), however only signature 16 mutations exhibit a higher preference for active domains (Supplementary Fig. 6e,h). Therefore, studying the distribution of mutations induced by individual signatures might help to distinguish the molecular underpinnings of different mutational processes.

## Unique folding of inactive X chromosome shapes the distribution of somatic mutations

Lastly, to investigate the impact of differential chromatin folding on somatic mutation distribution, we focused on X-chromosome mutations, as the inactive and active X-chromosomes exhibit distinct, well-studied folding structures that differ in males and females. Active X-chromosome (Xa), similar to autosomal chromosomes, is organized into discrete chromatin folding domains. Strikingly, inactive X-chromosome (Xi) is devoid of TADs; instead Xi is structured into larger repressive domains (Jégu et al., 2017). We utilized allele-specific X-chromosome chromatin contact maps (Darrow et al., 2016) to understand the somatic mutation distributions between female and male patients on the X chromosome. Notably, female X-chromosome mutation distribution is overall flatter compared to the mutations in male X-chromosomes (Fig. 6g, Supplementary Fig. 7a), with male X-chromosome somatic mutation distribution corresponding to TADs observed within Xa, similar to mutation distribution patterns in autosomal chromosomes (Fig. 6a). Strikingly, subclonal mutations on the X chromosome exhibit a flatter distribution compared to clonal mutations in female patients, which implies that Xi acquires more mutations during tumor evolution than the Xa (Fig. 6b). We did not observe any difference between the distribution of clonal and subclonal mutations in autosomal chromosomes (i.e. chromosome 7, Supplementary Fig. 7b), indicating this difference is specific to the X chromosome. Hypermutation of Xi has been reported in certain cancer types (Jäger et al., 2013). We sought to identify which cancer types exhibit higher mutation rate in X-chromosome in females. The biggest difference between male and female X-chromosome mutation load is observed in chronic lymphocytic leukemia samples. Female X-mutation load is significantly higher even compared to copy-number corrected male samples (p-value < 10^-6^; Fig. 6c-d). We checked whether males with X-chromosome polyploidy exhibit a similar mutation load to females but found no significant gain in mutation load from male polyploidy samples, implying the observed increase is predominantly due to inactive X-chromosome (Supplementary Fig. 7c). Brain cancers such as lower grade glioma (Fig. 6e), glioblastoma, pediatric brain tumors (Supplementary Fig. 7d-e) or thyroid adenocarcinoma and kidney renal cell carcinoma (Supplementary Fig. 7f-g) also present significantly higher X-chromosome mutation load in females compared to males. X-chromosome copy number and the expression of X-inactive-specific non-coding RNA (XIST) is significantly correlated with the X-chromosome mutation load predominantly in breast cancer samples (Supplementary Fig. 7h-i). Notably, certain cancer types do not exhibit a significant X-chromosome mutation load difference between males and females, such as lung adenocarcinoma (Fig. 6f). A subset of lung adenocarcinoma females (12 samples) shows hypermutation on X-chromosome but another subset (8 samples) contains similar level of X-chromosome mutations as males (17 samples) (Fig. 6f). We investigated the factors contributing to this variance and identified that signature 13 (APOBEC-related) occurs more frequently in females with lower X-chromosome mutations ratio. Signature 13 enrichment is negatively correlated with the X-chromosome hypermutation levels in female lung adenocarcinoma samples (r^2^: −0.42; Fig. 6g). This negative correlation between X-chromosome mutation ratio and APOBEC-related signature enrichments is observed in all female tumor samples across different cancer types (r^2^: −0.36; Supplementary Fig. 7j). As APOBEC-related mutations are enriched in active domains (Fig 5a), when APOBEC signatures are active, the ratio of Xi mutations is lowered. Taken together, our results demonstrate the importance of chromatin folding on the accumulation of somatic mutations in cancer cells. Differential chromatin conformations yield to distinct levels of somatic mutations accumulation, even though the underlying genomic sequence is identical and the activity of different mutational processes and chromatin configuration significantly contribute to the regional somatic mutation variability in cancer genomes.

**Figure 6.**
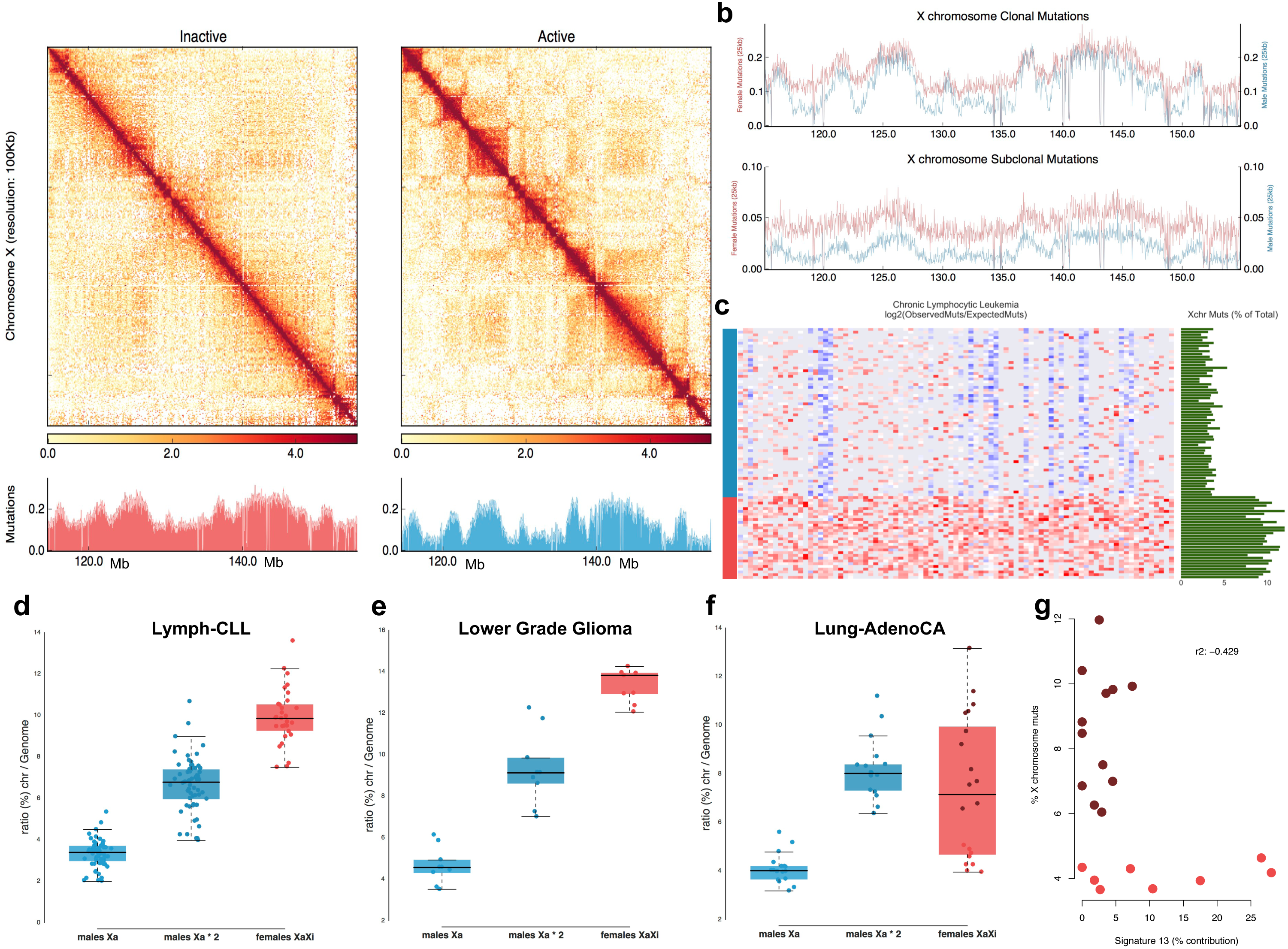
Unique folding of inactive X chromosome shapes the distribution of somatic mutations. **a)** Allele-specific Hi-C maps of inactive (Xi) and active (Xa) X-chromosome (between 115-155Mb) from human retina cells (RPE1 cell line). Below, histograms show the distributions of somatic mutation accumulation (in 25kb non-overlapping windows) per female (red) and male (blue) samples. Mutation numbers were normalized based on the number of samples. **b)** Mutation accumulation profiles are shown for the clonal (late) and subclonal (early) mutations along X chromosome (the represented part in Fig6a) in female (red) and male (blue) patients. Mutation numbers were normalized based on the sum of mutations in female and male cohort. **c)** Heatmap shows hypermutated TADs on X-chromosome in chronic lymphocytic leukemia cohort (TADs defined from non-phased Hi-C data). Bar charts show percent of X-chromosome mutations compared to all observed mutations in each sample (green) and normalized expression levels of Xist transcript (yellow). **d-f)** Distribution of X chromosome mutation load compare to total mutation burden for males (observed and copy-number corrected values) and females in **d)** chronic lymphocytic leukemia, e) brain lower grade glioma, **f)** lung adenocarcinoma cohorts. **g)** Negative correlation between APOBEC-related mutation signature (signature 13) and X-chromosome mutation accumulation in lung adenocarcinoma female patients.

## Discussion

Through integration of whole genome somatic mutation data and spatial genome organization, we have found that the activity of various mutational processes cause distinct mutation distributions across chromatin folding domains. Our results highlight that somatic mutation accumulation is correlated with genome organization. This differential distribution of somatic mutations can be observed in precancerous lesions such as in dysplastic Barrett’s esophagus (Supplementary Fig. 2d), implying that interaction of chromatin folding and mutagenic exposures begin to sculpt the mutation landscape early in carcinogenesis. Identifying the important long tail of driver cancer mutations is a challenge in cancer genomics (Martincorena and Campbell, 2015) and algorithms modeling the background mutation rate should take topological chromatin organization into account. Distribution of different mutational processes could point to underlying aetiologies of particular mutagens, such as signature 12 and signature 16 both frequently observed in liver cancers and exhibiting transcriptional bias. However, the distribution of mutations generated by these two signatures differentiate in terms of active versus inactive TAD enrichments. This could point a mechanistic difference between the aetiologies of these two signatures despite their strong association with liver cancer and strand bias similarities. In addition, when multiple mutational processes are operative in a tumor, the resultant mutation distribution is related to the rate and timing of each mutagenic factor. For example, MSI samples with high mutational signature 17 contribution deviate from flat-MSI distribution or induction of APOBEC mutagenesis due to tobacco smoking changes the initial mutational distributions. Shifts in mutational processes during later stages of somatic evolution of cancers could contribute to tumor heterogeneity as different genomic regions accumulate mutations based on the active processes. Focusing on the mutation distribution patterns along TADs could help to identify the biases generated by different mutational process over time and augments our abilities to elucidate the underlying aetiologies for specific processes – giving important insights into potential avenues to explore for therapeutic intervention.

The impact of chromatin folding is highlighted in female X-chromosome mutation distribution where there is marked interaction of mutation distribution with the distinct topology of inactive X-chromosomes. Subclonal mutation rate on Xi is higher compared to active X chromosome in female tumors, especially in cancer types where mutational processes preferentially target the inactive chromatin domains. The Xi observations could be illustrative of leveraging insights of mutational processes and chromatin folding for potential clinical benefit. Hypermutation is linked to immunotherapy response rate likely due to higher neoantigen production (Yarchoan et al., 2017), and activation of Xi is reported in human breast cancer models (Chaligné et al., 2015). Re-activation of Xi chromosome could potentially lead to higher neoantigen production and better response rate to immunotherapy in female patients and perhaps could be exploited clinically. More broadly, thorough understanding of chromatin folding and pervasive mutation processes may present opportunities. Inhibiting repressive epigenetic enzymes such as EZH2 might open up similar possibilities for modulating immunotherapy response rate in various cancer types as inactive domains with higher mutation load could lead to enhanced neoantigens expression upon EZH2 inhibition. Of note, 23% of coding mutations across the data set of 3000 genomes are in repressed chromatin domains, and thus may be potentially available to be exploited as neoantigens should efficient strategies for chromatin state modulation be determined.

Overall, chromatin conformation contributes to the variable accumulation of somatic mutations between and within human cancer genomes. Whether germline mutation distribution and genetic variations between different population groups or somatic mutation accumulation throughout aging process are correlated with the spatial genome organization remains an interesting open question for the future. Overall, investigating the three-dimensional genome architecture in human cancers will advance our understanding of mutational processes and DNA repair activity – ultimately painting a more informative, nuanced picture of this fundamental disease of the genome.

## Acknowledgements

We thank the patients and their families for contributing to this study. We thank Zeynep Coban Akdemir, Emily Keung, Paz Polak, Tony Gutschner for their critical readings of this manuscript. We also would like to thank the discussion of Xingzhi Song, John Zhang. Authors are grateful to all ICGC-PCAWG subgroup participants for generating uniformly analyzed mutation calls, mutational signature and mutation timing dataset. This work was supported by a Cancer Prevention Research Institute of Texas award (R1205), the Welch Foundation’s Robert A. Welch Distinguished Chair Award (G-0040 to P.A.F) and an US National Institute of Health Intramural Research Program Project Z1AES103266 (to D.A.G).

## Methods

### Data sources

We downloaded XR-seq data from GEO82213 and DNA lesions from GEO94434 et al. and replication-timing data from the ENCODE project data portal. Inactive X-chromosome and keratinocyte Hi-C interaction data were obtained from (Darrow et al., 2016), (Rao et al., 2014), respectively.

We obtained short tandem-repeat (STR) regions for the hg19 human genome assembly from gymreklab.github.io/software.html (Gymrek et al., 2017).

### TAD Annotation

We annotated TADs as described earlier (Akdemir et al., 2017). In brief, we used an entropy based approach (epilogos) to calculate occurrence of each chromatin state enrichment for a given genomic regions across all Roadmap Epigenome Consortia profiled cell types (compbio.mit.edu/epilogos/) (Roadmap Epigenomics Consortium et al., 2015). We calculated the ratio of a TAD genomic space covered by each chromatin state, divided by the length of the TAD, and generated a normalized matrix where columns are TADs and rows are each 15 chromatin states extensively studied by the Roadmap Epigenome Consortia. We applied hierarchical clustering to rows to identify similar chromatin states and k-means clustering to columns to group TADs containing similar epigenetic modifications. We decided on k=5 clusters as previous chromatin studies reveal 5 distinct epigenetically modified chromosomal domains and k=5 corresponded to better visually discernible domains.

### Mutation Calls and Slope Calculation

We utilized the International Cancer Genome Consortia’s pan-cancer (PCAWG) consensus mutation calls for the samples denoted as ICGC in Supplementary table 1 (synapse.org/#!Synapse:syn7118450). Whole genome sequencing data for other samples downloaded from EGA. We used Mutect (version 1.1.756) (Cibulskis et al., 2013) for calling mutations in these samples by using the default parameters. Identified mutations were filtered if the mutation is supported by at least 4 reads (t_alt_count > 4) in tumor sample and no reads in control (n_alt_count = 0). For ICGC samples, the active signatures in each sample were obtained from the PCAWG mutational signatures working group (synapse.org/#!Synapse:syn12025148) (Alexandrov et al., 2018). We used deconstructSigs (Rosenthal et al., 2016) to delineate the contribution of mutational signatures in a specific sample. MSI or PolE-deficiency information for colorectal, gastric and esophageal adenocarcinoma samples were obtained from TCGA Colorectal study (Cancer Genome Atlas Network, 2012), (Wang et al., 2014), (Secrier et al., 2016), respectively. Slope of samples calculated as the ratio of mutation burden in transcriptionally-inactive domains (low) versus transcriptionally-active (low-active) domains. We binned the mutations in 25 Kb non-overlapping windows along the genome. Mutation profiles were generated for each sample between transcriptionally-inactive (low) and –active (low-active) domains. Next, we calculated the total number of mutations, sum of NER-activity signal and replication timing within each TAD, and normalized the obtained sums with the given TAD’s length. For calculating enrichment of individual signature’s contribution to distinct domains, we plotted the activity of a specific mutational signature and the slope of mutations in each sample.

### APOBEC enrichment and Kateagis-like cluster analysis

APOBEC enrichments were calculated using a published method (Chan et al., 2015). Not APOBEC-enriched group represents samples with no statistically significant enrichment with the stringent tCa motif of APOBEC mutagenesis (tCaàtTa and tCaàtGa). APOBEC-enriched samples include both “A3A-like” and “A3B-like” groups of tumors. In order to remove a potential confounding factor, we excluded melanoma samples in this analysis because UV-mediated mutation signature resembles APOBEC signatures.

Kataegis-like G- or C-strand-coordinated mutation clusters were identified across PCAWG dataset (synapse.org/#!Synapse:syn7437313) as described in (Chan et al., 2015). Briefly, groups of closely spaced mutations in either C or in G of the top (sequenced) strand were identified, such that any pair of adjacent mutations within each group was separated by less than 10 kb and with p-value calculated based on mutation density in a sample of 10^-4^ or less. Shuffled clustered mutations are generated by randomly assigning clusters with keeping the number of boundaries per chromosome constant. Shuffling performed for 10,000 times for the clustered mutation set. We computed cumulative distribution of expected overlaps, z-scores and p-values were calculated based on observed number and obtained distribution from bootstrapping.

HBV, HCV and EBV annotations for liver and gastric cancers were obtained from Fujimoto et al., 2016, Wang et al., 2014, respectively.

### Mutation timing

Mutations for ICGC samples classified as clonal (early, late or NA) and subclonal by PCAWG tumor evolution and heterogeneity working group (synapse.org/#!Synapse:syn8532425) (Gerstung et al., 2017). This approach uses a hierarchical model to calculate timing parameters based on copy number and variant allele frequency data. If the mutation clonal frequency is equal to 1, those mutations are classified as clonal, whereas frequency lower than one determined as subclonal. We used the same method to identify mutation timing in esophageal MSI samples. (Code to determine timing of the mutations is available at github.com/gerstung-lab/MutationTime.R.)

### Cell lines

Two DNA-repair proficient (SW480, CaCo2) and two DNA mismatch-repair deficient (LoVo, DLD1) colon cancer cell lines were obtained from the American Type Culture Collection. Stocks were stored in liquid nitrogen. These cell lines were authenticated by morphologic inspection and short tandem repeat profiling. Mycoplasma testing was also performed in 2017 (A. Inoue). Cells were cultured in Dulbecco’s modified Eagle’s medium containing 10% fetal bovine serum and 1% Penicillin-Streptomycin at 37°C in a humidified incubator with 5% CO_2_.

We obtained whole-exome sequencing-based mutation calls for aforementioned colon cell lines from (Mouradov et al., 2014), Supplementary table 2.

### Hi-C experiment and processing

Hi-C was performed using the *in situ* Hi-C protocol as previously described in (Rao et al., 2014), using 2-5 million cells per experiment digested with the MboI restriction enzyme. Hi-C libraries were sequenced on a HiSeq 4000. Reads were aligned to the hg19 reference genome using BWA-MEM (Li, 2013), and PCR duplicates were removed with Picard. Hi-C interaction matrices were generated using in house pipelines, and matrices were normalized using the iterative correction method (Imakaev et al., 2012). HiCPlotter for visualizing normalized Hi-C interaction maps (Akdemir and Chin, 2015). In order to compare TAD boundary strength, we used quantile normalization to normalize different Hi-C data together.

